# Common properties of visually-guided saccadic behavior and bottom-up attention in marmoset, macaque, and human

**DOI:** 10.1101/2020.05.27.120428

**Authors:** Chih-Yang Chen, Denis Matrov, Richard Veale, Hirotaka Onoe, Masatoshi Yoshida, Kenichiro Miura, Tadashi Isa

## Abstract

The saccade is a stereotypic behavior whose investigation improves our understanding of how primate brains implement precise motor control. Furthermore, saccades offer an important window into the cognitive and attentional state of the brain. Historically, saccade studies have largely relied on macaque. However, the cortical network giving rise to the saccadic command is difficult to study in macaque because relevant cortical areas lie in sulci and are difficult to access. Recently, a New World monkey – the marmoset – has garnered attention as an attractive alternative to macaque because of its smooth cortical surface, its smaller body, and its amenability to transgenic technology. However, adoption of marmoset for oculomotor research has been limited due to a lack of in-depth descriptions of marmoset saccade kinematics and their ability to perform psychophysical and cognitive tasks. Here, we directly compare free-viewing and visually-guided behavior of marmoset, macaque, and human engaged in identical tasks under similar conditions. In video free-viewing task, all species exhibited qualitatively similar saccade kinematics including saccade main sequence up to 25° in amplitude. Furthermore, the conventional bottom-up saliency model predicted gaze targets at similar rates for all species. We further verified their visually-guided behavior by training them with step and gap saccade tasks. All species showed similar gap effect and express saccades in the gap paradigm. Our results suggest that the three species have similar natural and task-guided visuomotor behavior. The marmoset can be trained on saccadic tasks and thus can serve as a model for oculomotor, attention, and cognitive research.

**New & noteworthy:** We directly compared the results of video free-viewing task and visually-guided saccade tasks (step and gap) among three different species: the marmoset, macaque and human. We found that all species exhibit qualitatively similar saccadic behavior and bottom-up saliency albeit with small differences. Our results suggest that the marmoset possesses similar neural mechanisms to macaque and human for saccadic control, and it is an appropriate model animal to study neural mechanisms for active vision and attention.

## Introduction

Humans, like other primates, rely heavily on vision for daily life. Only the center of our retina has high visual resolution, so we must constantly move our eyes to collect information about the world in an effective manner. Three to four times per second our brain sends a signal to point our eyes to a new location using a fast, ballistic rotation of the eyeballs known as a saccade (Yarbus 1967).

Our understanding of the neural mechanisms underlying the human visual and oculomotor systems and the interaction between them are based largely on research in macaque (e.g., Felleman and Van Essen 1991; Fuchs 1967; Knöll et al. 2018; Markov et al. 2014; Mustari 2017). Indeed, macaque has a visual brain remarkably similar to human and has been the major source of our understanding on how the visual cortex functions (Hubel and Wiesel 1969; Vanni et al. 2020), as well as how eye movements are targeted and timed (Krauzlis 2005; May 2006; Moschovakis et al. 1996; Schall 2015; Shinoda et al. 2019). The interaction between the two systems, including how a visual target is selected (Medendorp et al. 2011; Schall et al. 2011; Schiller and Tehovnik 2005) have also been investigated. Inspired by the physiological research, computational models have emerged to predict how primates select where to look when freely viewing pictures or videos (Borji et al. 2013; Itti and Koch 2000; Kümmerer et al. 2018; Veale et al. 2017). These models mimic the visual hierarchical brain to filter visual features such as luminance, spatial frequency, orientation, and motion to produce an estimate of the distribution of “saliency” or “conspicuity” across the visual field to predict probability of saccade targets. Despite the excellent progress made in saccade research, we are still unable to exhaustively explain the factors that contribute to a given saccade to a given target at a given time. To complete our understanding, further research is necessary into the interactions of the myriad of cortical and subcortical areas involved in the saccade process.

Recently, the smaller and more manageable New World monkey, *Callithrix jacchus* (the common marmoset) has been proposed as an attractive alternative to the macaque for neuroscience research (Homman-Ludiye and Bourne 2017; Mitchell and Leopold 2015; Solomon and Rosa 2014). Besides the increasing availability of transgenic animals (Belmonte et al. 2015; Park et al. 2016; Sasaki et al. 2009; Tomioka et al. 2017b, 2017a), one important advantage is that the marmoset has a lissencephalic (smooth) brain, facilitating access to multiple cortical areas simultaneously (Isa 2017; Tia et al. 2017; Umeda et al. 2019). Recently, efforts have been made to demonstrate that the marmoset is intelligent enough to be trained on oculomotor tasks and compare its saccadic kinematics to macaque (Mitchell et al. 2014). However, experimental details including inconsistent eye movement recording devices and sampling frequencies have made the comparisons difficult to interpret – thus motivating the current report.

Besides basic saccadic kinematics, one phenomenon that has been widely studied in oculomotor research is the “express saccade”: a special type of saccade exhibiting ultra-short response time in response to a visual target appearing (Fischer and Boch 1983; Fischer and Ramsperger 1984). Normally, a saccade responding to a visual target involves several brain areas (Isa 2002; Krauzlis 2005), and the process from the onset of a visual stimulus to the onset of physical eye movement takes between 150 and 200 ms. However, if a blank screen is inserted for a short “gap period” before the appearance of the visual target, the time to respond to the visual target can be facilitated (this is called the “gap effect”). Sometimes, this happens so often that the distribution of saccadic reaction times forms a clear bimodal distribution, with the first peak (representing express saccades) at around 70 to 80 ms in macaque. Insertion of the “gap period” in the visually guided saccade paradigm has become so popular that it has received its own name: the “gap saccade task”. Recently, one study described training marmosets in the gap saccade task, but only observed the gap effect, not the appearance of a full bimodal distribution (Johnston et al. 2018). This difference from macaques has made it ambiguous whether marmosets are appropriate for studying express saccades.

In the current study, we will directly compare between species both saccade kinematics and saliency map prediction during an untrained video free-viewing task, as well as comparing oculomotor response time during trained visually-guided step and gap saccade tasks. We will do this for three species: marmoset, macaque, and human. Our hypothesis is that marmoset, macaque, and humans all share similar saccade kinematics and basic saliency-driven aspects of looking behavior. We compare the three species under nearly identical experimental and recording conditions for each task. We conclude that the visual and oculomotor behavior of marmoset share common mechanisms with macaque and human. This indicates that study of the brain mechanisms responsible for marmoset visual and oculomotor behavior can be generalized to macaque and also human.

## Materials and Methods

### Subjects

Three marmosets, two macaques and four humans were subjects in this study. All animal experimental protocols for macaques and marmosets were in accordance with the guidelines for animal experiments approved by Animal Ethical Committee in Kyoto University, Japan. All human experiments were conducted in accordance with the Declaration of Helsinki and approved by the Kyoto University Graduate School and Faculty of Medicine, Ethics Committee. Informed consent was obtained from each subject after a full explanation of the study procedures. Anonymity was preserved for all participants. We recruited 2 human subjects (Human S1 and S2, both male) for this study. Two other subjects were the first and the second author of this study (Human C and Human D, both male).

### Animal preparation

We collected behavioral data from 3 adult marmosets (*Callithrix jacchus*). Marmoset A (3 years 6 months, 325 g) and Marmoset B (3 years 3 months, 314 g) were male weighing 320 and 310 g, respectively. Marmoset C (3 years 3 months, 301 g) was female and weighed 330 g. All marmosets first underwent 2 to 3 weeks of marmoset chair (MMC-3505, O’HARA & CO.) adaptation before any surgery. After the marmosets could quietly sit in the chair and lick the periodically administered reward (described below) for more than 20 min, they were surgically prepared for head-fixed eye movement measurement. Under isoflurane anesthesia using a facemask modified from “Johnston et al. 2018”, we attached one head-holder for each marmoset. The head-holders were either custom made by our machine shop with PMMA or 3D-printed with bio-compatible material (Form2, Formlabs). Location of the implant was predetermined based on CT and MRI scans for each marmoset. The bottom of the head-holder was first attached to the skull with Bistite II (Tokuyama Dental) and further supported with Unifast II dental resin (GC Corporation).

We also collected behavioral data from 2 macaques (*Macaca fuscata*). Monkey O (7 years old, 6.1 Kg) was female and Monkey U (4 years old, 6.2 Kg) was male. Under isoflurane anesthesia with intubation, each macaque was implanted with head-holders made of POM surrounded with skull screws as anchors. The head-holder and the skull screws were further supported with Unifast II dental resin (GC Corporation).

### Eye tracking method

For marmosets and macaques, we used an Eyelink 1000 Plus (SR Research, Ontario, Canada) to track their eye. For humans, an Eyelink 1000 (SR Research, Ontario, Canada) system was used. All the subjects were tracked with 500 Hz sampling rate on their right eye.

### Calibrating eye position

In order to gain better control over the eye position during the tasks, we used a two-step calibration for marmosets. We first applied a rough calibration using marmoset face images similar to “Mitchell et al. 2014”. Briefly, we presented marmoset face images (8 images each trial, image size ranging from 4 to 20.25 deg^2^) in random positions on the screen. The positions were controlled so that the images never overlapped with one another. The marmosets were allowed to look at the images for 10 second each trial. After collecting 6 trials, the experimenters used a custom user interface to inspect simultaneously for all 6 trials where the images were presented on that calibration trial along with the corresponding collected gaze positions for that trial. With this user interface, we were able to adjust the offset and the gain of the eye positions from the 6 trials simultaneously so that the eye positions overlaid with the location of the images. We usually did this calibration only once after the marmosets were head-fixed and sat in front of the monitor. After the first calibration, the offset and gain were fed into our eye controller system. To further refine the calibration, the marmosets performed 48 trials of visually-guided step saccade task (described below). For each trial, we took the averaged eye position 50 ms before the marmosets got their reward and calculated the offset and gain again using simple linear regression. We then corrected our system using the new offset and gain. After the eye position was calibrated the first day, we performed only 24 trials of visually-guided saccade task to validate and make minor changes on the gain and offset in later days. The same calibration procedure was used for macaques and humans.

### Behavioral tasks

#### Video free-viewing task

To study saccade kinematics and gaze behavior under more natural conditions, we presented the same video stimuli as “Yoshida et al. 2012 (the CRCNS-ORIG database – Itti and Carmi 2009)” to all our subjects across species (Fig. 1). Videos from the database were displayed in their native pixel size (558 - 640 pixels in width, 394 - 480 pixels in height) over a gray background (80.81 cd/m^2^) to avoid any potential truncation at the edge of the videos. The length of video ranged from roughly 4 - 94 sec. Each marmoset sat in the marmoset chair with its head fixed using the head-holder and facing a 1280 X 1024 pixels LCD monitor with 375 mm in horizontal length. The viewing distance was 40 cm, resulting in a 614 X 448-pixel video to have 24.1 X 17.6-deg viewing angle (Fig. 1A, left panel). The marmosets received a drop of reward consisted with banana pudding (Kewpie Corp.) and banana flavored Mei Balance enteral supplement (Meiji Holdings Co.) every second to keep them awake and paying attention to the video. The reward delivery was accomplished by using a custom-made syringe pump (O’HARA & CO.) with high precision and torque designed for viscus fluid. Since we did not restrict their gaze location to be within the video range, the animals were free to look at any part of the setup that their oculomotor range would allow, including the edges of the monitor or gray background around the video. We collected 124 to 150 trials (one random video presentation each) of looking data for each marmoset.

**Figure 1.**
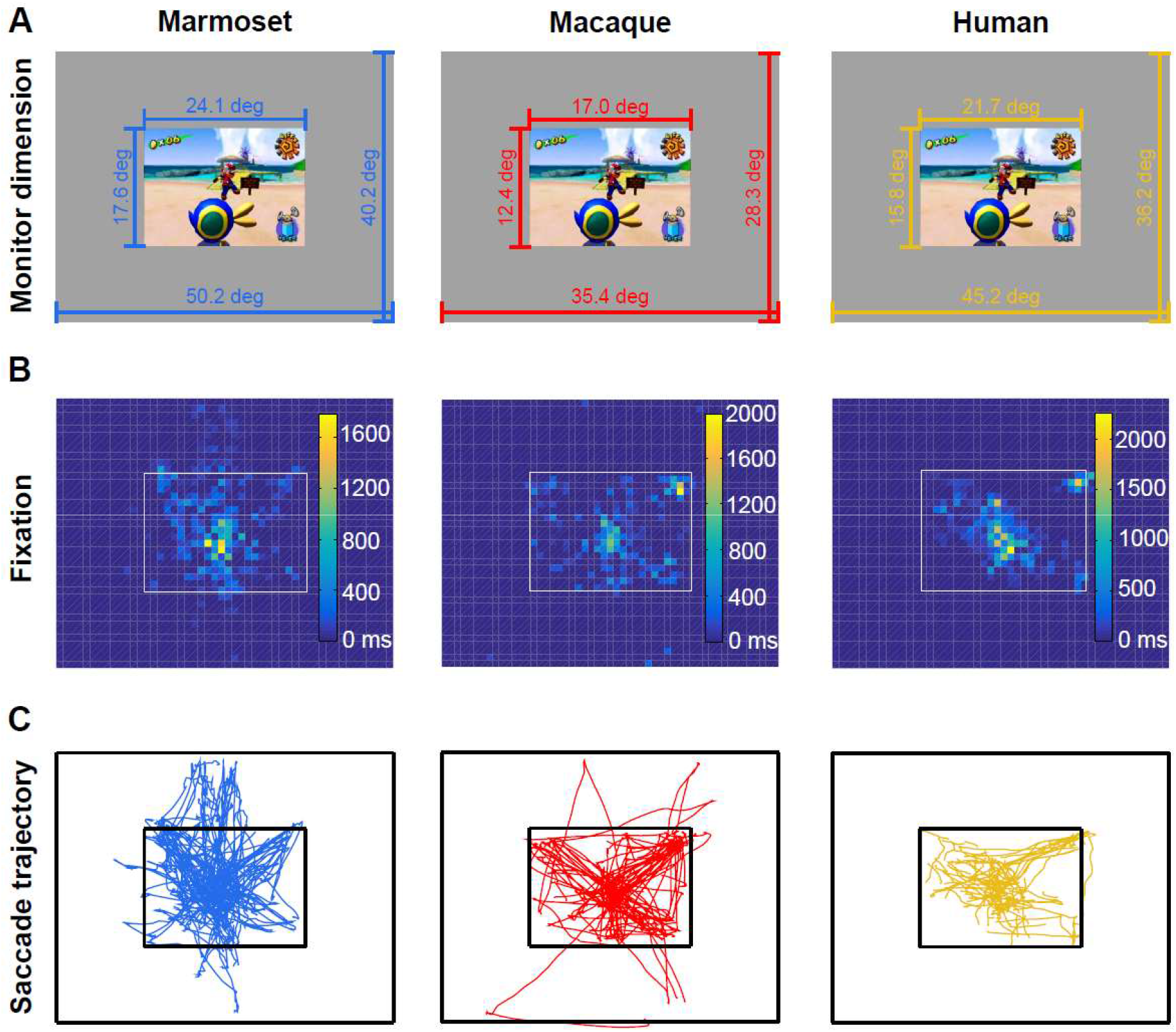
Example video free-viewing task session from the three species. **(A)** Monitor and video dimension used for each species. The gray background in each column indicates the monitor area. Dimension are labeled with horizontal and vertical visual angles calculated from the relative distance of the monitor to the eye position of each species in their setups. The extent of the video stimulus embedded in the monitor background is also labeled in degrees visual angle. **(B)** Accumulated fixation time across the same example video. We plot the total fixation time across the selected example video within the same pixel area from one representative subject of each species in each column. From total fixation time for each representative subject, the marmoset spent 86%, macaque spent 80%, and human spent 96% of fixation time within the video range of this example video. **(C)** Saccade trajectories across the same example video. We plot saccades made by the three representative subjects separately in each column.

The macaques were also head-fixed and sat in their own macaque chair facing front to the monitor. The monitor was an LCD monitor with 1920 X 1080 pixels and 600 mm in horizontal length. We covered the surround of the monitor with black paper and eventually made the visible area to be 1280 X 1024 pixels at the center of the monitor. The viewing distance was 60 cm for macaque setup, resulting in the same 614 X 448-pixel video to have 17.0 X 12.4-deg viewing angle (Fig. 1A, middle panel). The gray background behind the video was 103.7 cd/m^2^. During video free-viewing task, a drop of water reward was also delivered every second with a custom-made solenoid. As with marmosets, gaze location was not restricted to the video area. We collected 99 trials (again, one video each) for Monkey O and 187 trials for Monkey U.

In the human setup, we used a monitor with similar size and pixel dimension to the monitor used in marmoset experiments. The viewing distance was 45 cm, resulting in the same 614 X 448-pixel video subtending 21.7 X 15.8-deg visual angle (Fig. 1A, right panel). We used a head support (SR Research, Ontario, Canada) for restraining the head of our subjects. We instructed the subjects to freely look at the videos. The gray background behind the video was 220.2 cd/m^2^. We collected 126 to 143 trials for each human subject.

We endeavored to maintain identical conditions among all three species. However, due to geometry limitations of our experimental spaces, the distance from the subject to the stimulus presentation monitor varied, resulting in minor differences in viewing angle between the three species (marmoset / macaque: ~1.42 times; human / macaque: ~1.28 times; marmoset / human: 1.11 times larger viewing angle). In all cases, videos were played in their original size, which is relatively small (Fig. 1A) in all experimental setups, all videos were well within the oculomotor range of all species and subjects (Fig. 1B-C). Another purpose for playing the videos in their original size was to preserve the quality of the video frames and facilitate the later bottom-up saliency model calculation (Fig. 3).

**Figure 2.**
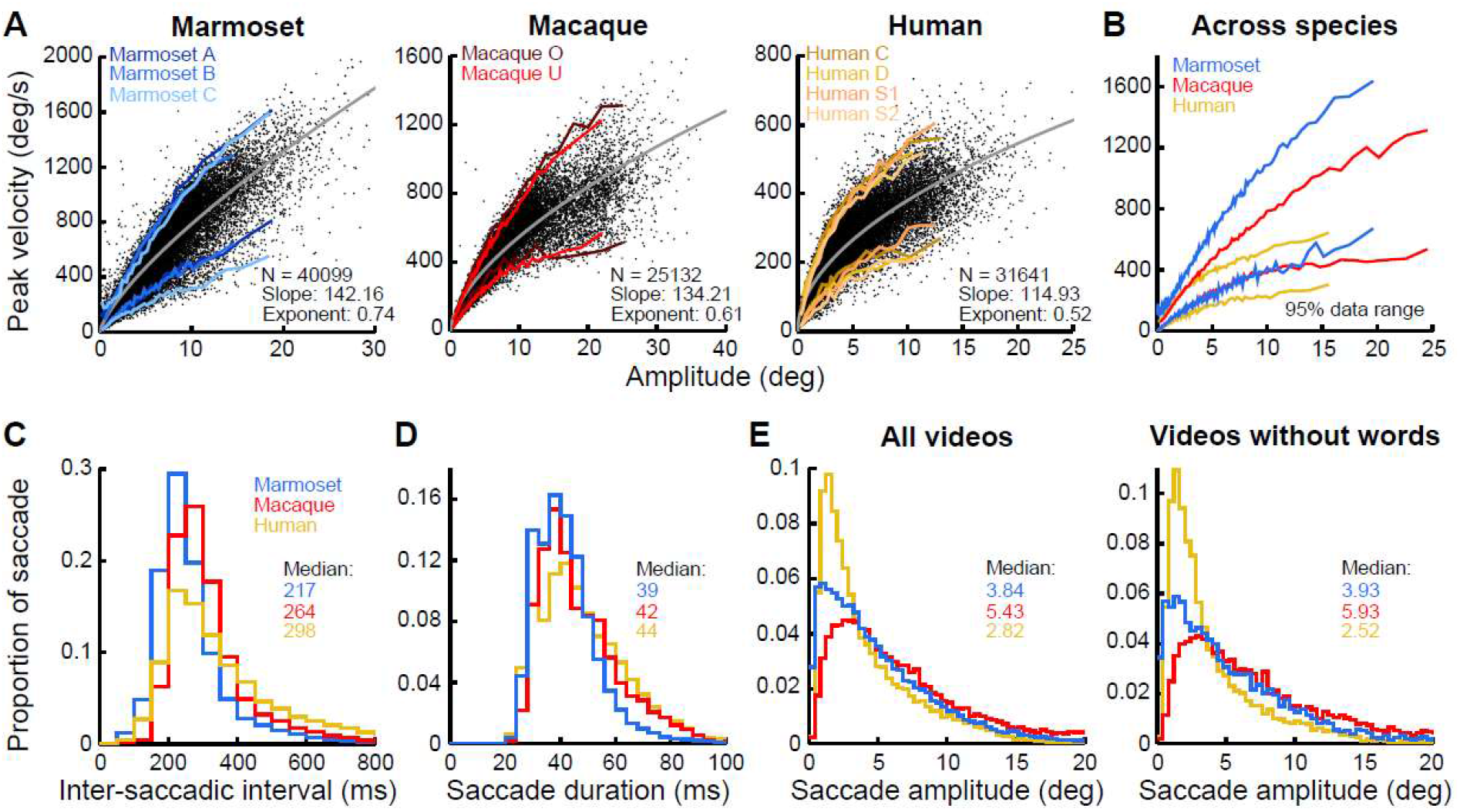
Saccade kinematics collected from video free-viewing task. Data from all subjects separated on a per-species basis. **(A)** Main sequence of the three species. The saccade amplitude against the peak velocity is plotted separately for each species. Black dots indicate all saccades collected from the task. N represents total number of saccades collected from each species. The gray smooth curve along the center is the curve fitted to the equation described in Materials and Methods for each species. The 95% data range boundaries are plotted around the dot clouds for each subject separated with color indication. **(B)** The 95% data range of the main sequence from each species. To facilitate comparison, the 95% data ranges of the main sequence from each species are re-calculated and plotted. **(C)** Inter-saccadic interval for each species from video free-viewing task. **(D)** Saccade duration for each species from video free-viewing task. **(E)** Saccade amplitude for each species either from looking at all the videos (left) or only the videos without words (right). The median of the data distribution for each species in (C-E) is indicated in the intersect.

**Figure 3.**
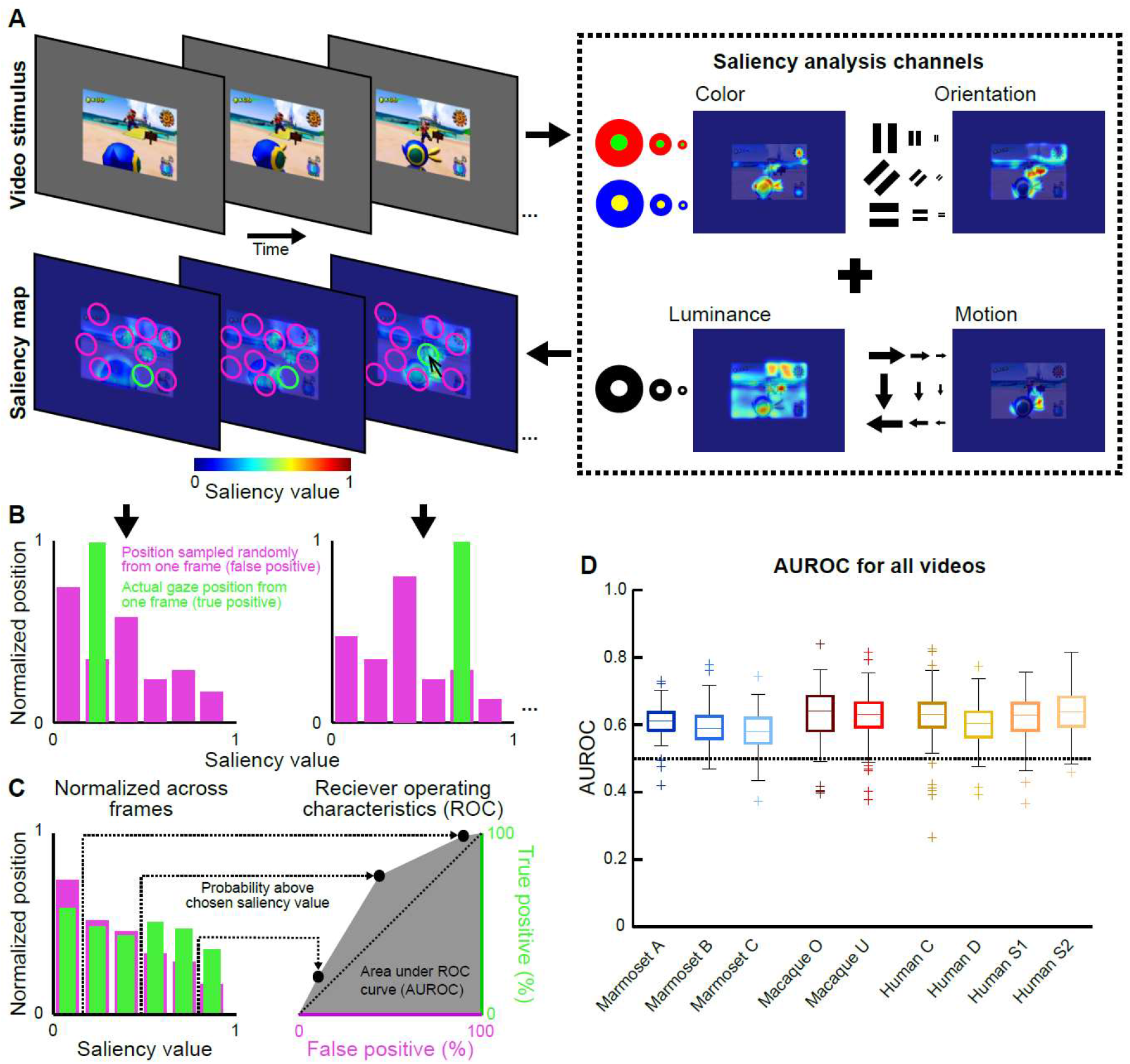
Implementation of bottom-up saliency model and the result in predicting gaze position for all subjects. **(A)** Frames from our video stimuli (top) and overlaid saliency map model results (bottom). The location of a subject’s gaze (true positives – green circle in each frame) during a given video frame is compared against other locations that the subject could have, but did not, look (magenta circles) drawn from the subject’s prior probability distribution of looking. **(B)** For each frame in each trial, an instantaneous distribution of the saliency values for the true target of looking (green bar) and false target (magenta bars). **(C)** We combine the true targets and false targets from all saccades in a trial into a single distribution (left) representing the saliency distribution of true (green) and false (magenta) targets. The predictivity of the model is then quantified as the area under the receiver operating characteristic ROC curve (AUROC) that is swept out by plotting true positive to false positive detection ratios at different sensitivity thresholds (shaded gray area – right). **(D)** Distribution of AUROC values for each subject computed separately for each trial (single viewing of a single video).

#### Step saccade task

A fixation spot consisting of a white circle (0.33 deg in diameter) surrounded by black arena (total 1 deg in diameter) was presented over a gray background at the center of the monitor (Fig. 4A). After the subject fixated for 200 ms within an invisible zone defined by a 2 deg radius circle centered at the fixation spot, the fixation spot jumped randomly to one of eight locations, all at the same visual eccentricity (6 deg, if not mentioned elsewhere) but separated by 45 deg in radial direction from right horizontal of the monitor (0, 45, 90, 135, 180, 225, 270, or 315 deg). The jumped fixation spot served as the peripheral saccade target. The subject was either instructed (for human) or trained via liquid reward (for macaque and marmoset) to make a saccade to the saccade target. Each species performed the same task with the only difference being the differences in the setups described above. We collected 630 trials for Human C, 669 trials for Human D, 607 trials for Human S1, 592 trials for Human S2, 800 trials for Macaque O, 1431 trials for Macaque U, 1049 trials for Marmoset A, 905 trials for Marmoset B, and 793 trials for Marmoset C.

**Figure 4.**
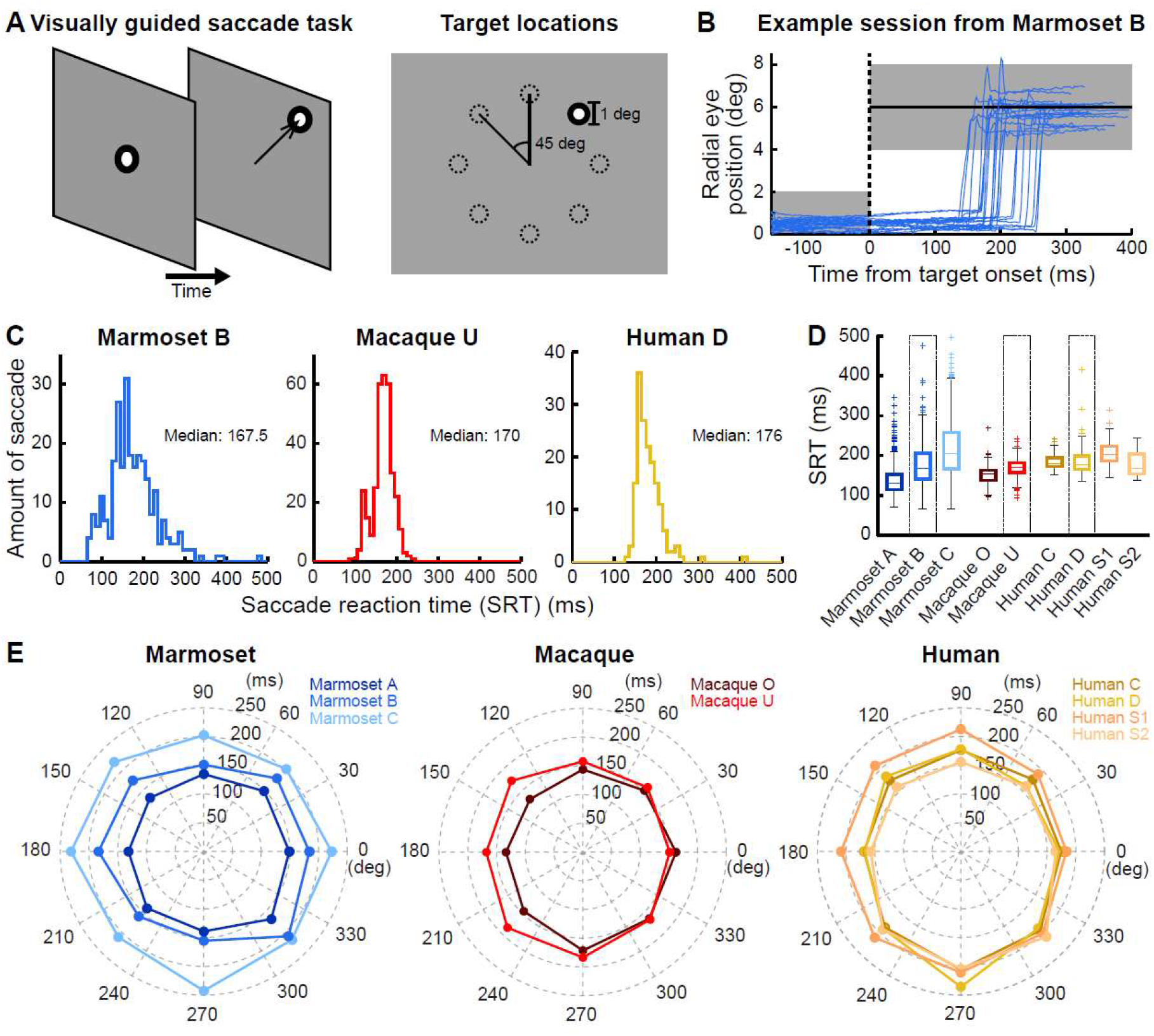
Performance in visually-guided step saccade task in the three species. **(A)** Diagram of visually-guided step saccade task. Gray diamond shape indicates monitor event on the left panel. The black arena with white dot indicates either fixation target or saccade target. The size and the potential saccade target locations are indicated in the right panel with dashed circles. The targets were at fixed radial eccentricity and separated by 45-degree radial angle. The targets appeared in random order. **(B)** Example session from a marmoset performing visually guided saccade. Time from target onset is plotted against the radial eye position from Marmoset B. Dashed line indicates target onset time. Gray shaded area before and after the dashed line indicates the eye window for both online gaze control and post-hoc analysis. **(C)** Example saccadic reaction time (SRT) histogram for each species. The example subjects from each species were chosen because the median of their SRT was similar. **(D)** Box plot for SRT in all subjects. Subjects are plotted in different colors. Middle mark in the box indicate the median of the SRT. The upper and lower bound of the box represent the 75th and 25th percentiles, respectively. The crosses (+) indicate outliers for each species. The example subjects we used for each species are indicated with a black box around their box plot. **(E)** Mean SRT for all subjects separated by target locations. The radial axis plots the SRT against the angular axis with target locations. The rightmost horizontal target is 0 degrees radial angle, according to convention. Marmosets, macaques, and humans are plotted from left to right.

#### Gap saccade task

The same fixation spot and target parameters used in the step saccade task were also used for the gap saccade task. After the subject fixated the fixation spot for 200 ms, the fixation spot was removed, leaving only the gray background visible. The subject had to maintain fixation within the invisible zone defined by a 2 deg radius circle around the position of the (now disappeared) fixation spot. After a fixed time for each session (gap period; either ~50, 100, 150, 200, or 250 ms, randomly chosen for each session), a peripheral saccade target was presented identical to the one used in step saccade task. Every session was terminated with 24 successful trials or when the total trial number (both successful and failed trials) reached 48. The reason for inter-session randomization for different gap periods was to facilitate training for marmosets. For fair comparison, all species performed exactly the same task in their own setup described above. We collected per gap period 610 trials for Human C, 624 trials for Human D, 629 trials for Human S1, 604 trials for Human S2, 505 trials for Macaque O, 726 trials for Macaque U, 539 trials for Marmoset A, 484 trials for Marmoset B, and 414 trials for Marmoset C.

### Saccade detection

We used the same previously established method in “Chen and Hafed 2013” to detect saccades in all experimental conditions (video free-viewing, step, and gap) for all species. Briefly: we first linearly interpolated the saccade data to 1 kHz. We then identified all time samples that exceeded 15 deg/s in radial eye velocity. We further refined the start and end points of saccades until eye acceleration reached 500 deg/s^2^ radius. We manually inspected each saccade to correct for false alarms or misses. Finally, blinks or noise artifacts were manually marked for later removal.

### Data analyses

#### Video free-viewing task, example trials

To plot the accumulated fixation time of the example video shown in Fig. 1B, we first selected only fixational gaze positions by removing all the detected saccades, blinks, and noise artifacts from the example trail. The remaining time periods (regardless of their content) were defined to be “fixations”. We then cut the area of the video into the same amount of small squared pixel areas (25.5 pixel on each side) for each species and we calculated the accumulated fixation time by accumulating the fixational gaze time within the small squared pixel areas from trial start to trial end. We also extended these areas out from the video area and cover the entire monitor range. Fig. 1C shows only those saccades detected from the single trial we used for the example.

#### Video free-viewing task, saccade kinematics

To characterize the main sequence (Bahill et al. 1975), we first fitted the amplitude versus the peak velocity of the saccades collected from video free-viewing task with the following equation for each subject:

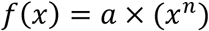

where *x* is saccade amplitude, and *f*(*x*) is saccade peak velocity. The fitted parameter *a* and *n* represent the slope and exponent of the fitted curve, respectively. The parameters *a* and *n* did not significantly differ within each species (Table 1). This is expected because the main sequence has been widely reported to be stereotypic within species. We then combined data for each species and fitted again using the same equation (Fig. 2A, B).

**Table 1.** Main sequence parameters for individual subjects. The subjects are grouped within species. a and n are the slope and exponent parameter for fitting, respectively. R^2^ represents the goodness-of-fit for the function. N represents the number of saccades observed for each subject.

|  | Marmoset |  |  | Macaque |  | Human |  |  |  |
| --- | --- | --- | --- | --- | --- | --- | --- | --- | --- |
|  | A | B | C | O | U | C | D | S1 | S2 |
| a | 141.74 | 149.99 | 125.12 | 168.63 | 118.13 | 132.87 | 114.18 | 104.20 | 100.51 |
| n | 0.77 | 0.73 | 0.76 | 0.53 | 0.66 | 0.47 | 0.50 | 0.59 | 0.55 |
| R<sup>2</sup> | 0.91 | 0.86 | 0.81 | 0.81 | 0.87 | 0.81 | 0.76 | 0.85 | 0.82 |
| N | 16225 | 12941 | 10933 | 8180 | 16952 | 9633 | 6839 | 7752 | 7417 |

We then calculated saccade duration, amplitude, and inter-saccadic interval for each species (Fig. 2C-E). Inter-saccadic interval was defined as the time between the end of a saccade and the beginning of the next saccade. Saccade amplitude was the Euclidian (straight-line) distance between the starting point and ending point of the saccade (i.e. not the trajectory integral). Saccade duration was the difference between the start and end time of the saccade.

#### Video free-viewing task, bottom-up saliency model implementation

To further study potential similarity of the bottom-up driven saccades across subjects, we also implemented the saliency map model developed by “Itti and Koch (2000)”, including channels for luminance, color, motion, and orientation (Fig. 3). The saliency map model was used to create a distribution of looking probabilities for each frame of our video free-viewing task. We used a library developed in-house (https://github.com/flyingfalling/salmap_rv) which allows easy parameterization of the saliency map model. Specifically, in our case, we first generated a gray background with raster R/G/B image similar to our monitor size (1280 X 1024 pixels) and fed it into the model. Within the gray background, we embedded the video stimuli at their original pixel size which was shown to all subjects (Fig. 3A). To convert pixel space to visual angle space for each species, we calculated the degrees visual angle per pixel (0.039242 for marmosets, 0.027672 for macaques, 0.035344 for humans) based on the behavioral setups described earlier and also used these numbers in the model to scale the video frames input to the saliency map model. Since the saliency map model can be parameterized in many ways, we briefly describe the model used to generate these results, and refer users to the configuration file for the model available in the github repository above, in models/chenetal2020marmo.params.

As a first effort, we implemented the computational “saliency map” model of “Itti and Koch 2000”, with an additional motion channel to capture perceived motion. The model separates each color video frame into four visual feature channels corresponding to (1) Luminance (2) Orientation (3) Color-Opponency and (4) Motion (see Fig. 3A, right, for an example of the output of these 4 feature channels). The saliency model detects each visual feature channel at multiple spatial frequency scales, and then subtracts low-frequency scales from high-frequency scales (a center-surround mechanism) to detect regions of local difference. Center-surround levels are combined and normalized within each feature channel, and then between all the different feature channels to produce a final, feature-agnostic saliency map, which is what we use to predict where a subject will look while viewing that video.

Our model computes luminance via the simplest method, as the average value of (red, green, blue) components. The model uses four different orientation angles (0, 45, 90, 135 degrees) computed using Gabor filters on a high-pass filter output of each scale of the luminance map, and four motion directions (0, 90, 180, 270) computed using Reichardt detectors on the luminance map. Additionally, we use two color opponency channels (Blue-Yellow and Red-Green), computed using the algorithm from “Walther and Koch 2006”. To produce a map at a given spatial frequency scale, we low-pass filter the original using a Gaussian kernel of the appropriate width.

We do not know *a priori* what the “best” spatial frequencies are for each species or subject, and determination of these values is outside the scope of this paper. We intend the saliency map model to serve as an approximation of the types of features that will draw bottom-up visual attention. To maintain fairness and to prevent unintended biases, we use the same model with the same spatial frequency tunings for all subjects and all species, with no optimization of parameters. This will allow us to directly compare how much basic visual features draw the attention of subjects from each species. The results reported in this paper are produced by a model that samples each channel at spatial frequencies of 8 Hz, 4 Hz, 2 Hz, 1 Hz, 1/2 Hz, 1/4 Hz, 1/8 Hz, and 1/16 Hz (cycles per degree visual angle, see table), with each level low-pass filtered from the preceding level using a Gaussian kernel with a width tuned to pass 90% of the specified frequency’s power unattenuated. The high-pass filter used for the orientation channel is computed simply by subtracting the high-frequency map from the adjacent lower-frequency map.

| Map Level | Spatial Frequency (in cycles/deg visual angle) |
| --- | --- |
| 0 | 8 Hz |
| 1 | 4 Hz |
| 2 | 2 Hz |
| 3 | 1 Hz |
| 4 | 1/2 Hz |
| 5 | 1/4 Hz |
| 6 | 1/8 Hz |
| 7 | 1/16 Hz |

The model computes local differences within each feature channel by computing the difference at 3 different “center” levels from 2 “surround” levels each. The surround levels are offset by 3 and 4 levels from the center levels. Thus, the center-surround difference levels we use are: **[1-4], [1-5], [2-5], [2-6], [3-6], [3-7]**.

After downsampling each local difference map to the output spatial scale (0.5 degree visual angle per pixel), the model normalizes the maps using the LMAX (local max) algorithm described in “Itti and Koch 2000”, and then takes the average within each feature channel sub-channel (specific motion direction, orientation angle, or color opponency channel) and normalizes the result again using LMAX. All sub-channels are then averaged again and normalized with LMAX, followed by a final average among all feature channels and a final LMAX normalization. This is the final map we use for predicting looking, with one additional caveat: we apply a temporal low-pass filter to the final saliency maps for each frame with a time constant of 100 ms. The purpose of this final low-pass filter is to smooth artifacts due to discretization in motion that may occur due to the frame-rate of the video or the method of pixel sampling. This temporal low-pass filtered version of the feature-agnostic saliency map is what we use to predict looking for each subject.

#### Video free-viewing task, looking behavior predication by saliency model

We compute the accuracy of the saliency map model for each trial for each subject. First, we construct a “null model”, which is set of saccade endpoints of the subject on every other trial except for the current trial. This set of “possible” looking positions are used to approximate the prior probability distribution of the subject’s looking, i.e. it takes into account any lateral or center biases the subject may have (Fig. 3A, magenta circles). This method has previously been described as the “shuffled AUC” in previous papers (Borji et al. 2013; Tatler et al. 2005; Zhang et al. 2008).

To quantify how much the saliency map model predicts where a subject will look, we use the Receiver Operating Characteristic (ROC), which is a comparison of the true-positive detection rate versus the false-positive detection rate of the model for a range of sensitivity thresholds.

For each saccade in the trial, we find the saliency distribution computed by the saliency map model for the video frame directly preceding the onset of the eye movement (Fig. 3, top-left). First, we determine the “true positive” rate by taking the average saliency value within a circle with radius 0.75 deg visual angle around the endpoint location of the eye movement (Fig. 3A, cyan circles). Then, the “false positive” rate is the mean saliency value within 1000 similar circles with centers randomly drawn from the subject’s prior looking distribution described above (Fig. 3A, magenta circles – for demonstration, only 10 circles are shown). The distribution of these saliency values can be visualized as a histogram as in Fig. 3B, where magenta represents the probability distribution of saliency values of the prior draws, and cyan represents the probability distribution of saliency value of the true target. Because there is only one true target for a single saccade/frame (i.e. where a subject actually looked), there will always only be a single cyan bar containing 100% of the density for the histogram we draw for a single saccade. We combine all saccade histograms from a trial into a single histogram (Fig. 3C). From this trial distribution, we compute the x-values of the ROC curve: the true positive (cyan) density above each threshold for thresholds in increments of 0.05 from [0,1] (Fig. 3C, left panel). The same is done to determine the y-values of the ROC curve: these are the false positive (magenta) density above each threshold. Each pair of values (true positive rate above threshold, false negative rate above threshold) specifies a point on the ROC graph, leading to 21 points for 21 different thresholds (in Fig. 3, only 3 are shown). We condense this curve into a single number by taking the integral area under the ROC curve (AUROC) using the trapezoid method (Fig. 3C, right panel). An AUROC of 0.5 indicates chance level, which means the saliency of the target of an eye movement is just as likely to be salient as any other place the animal could reasonably had looked at that time, given the subject’s prior distribution of looking locations. An AUROC above 0.5 indicates the target of looking was likely to have higher saliency than other places the subject would normally have looked.

#### Step and gap saccade task, saccade reaction time

We extract the saccade reaction time (SRT) in the visually-guided step and gap saccade tasks. First, we determined which saccade was the “response” saccade to the visual target in the task. We took the first saccade with saccade landing position less than 2 deg relative to the saccade target location after saccade target onset. We then traced backward in time and excluded trials if there was a microsaccade (saccades with <=1 deg during fixation period) before this saccade up until 100 ms preceding target onset. We excluded these trials because it has previously been shown that microsaccadic suppression may affect SRT (Hafed and Krauzlis 2010). After excluding trials with microsaccades, we further excluded trials in which the response saccade could have been a guess (anticipatory saccade). To facilitate fast training in the marmosets, we did not randomly vary the fixation time in the visually-guided step nor the gap saccade tasks. We also used a block design for the gap period – i.e. the gap period was the same for all trials within a given training session. Under these conditions, subjects could memorize the gap timing and make saccades anticipating the visual target appearing, even if they could not yet see the target. To exclude trials in which such “anticipatory saccades” were made, we first defined the potential anticipatory saccades to be the saccades that fit with the above criteria for SRT extraction but whose landing position was not within the 2 deg window around the saccade target (Fig. 5C, thin black line). Based on the distribution of timings of these “potential anticipatory saccades” for each species, we set thresholds for minimum SRT to be 65 ms for marmosets, 85 ms for macaque, and 100 ms for human. Any saccades with reaction times shorter than these thresholds were assumed to be anticipatory, and excluded from further analysis. Around 1.5% of the trials in humans, 2.5% of the trials in macaques, and 6% of the trials in marmosets had anticipatory saccades.

**Figure 5.**
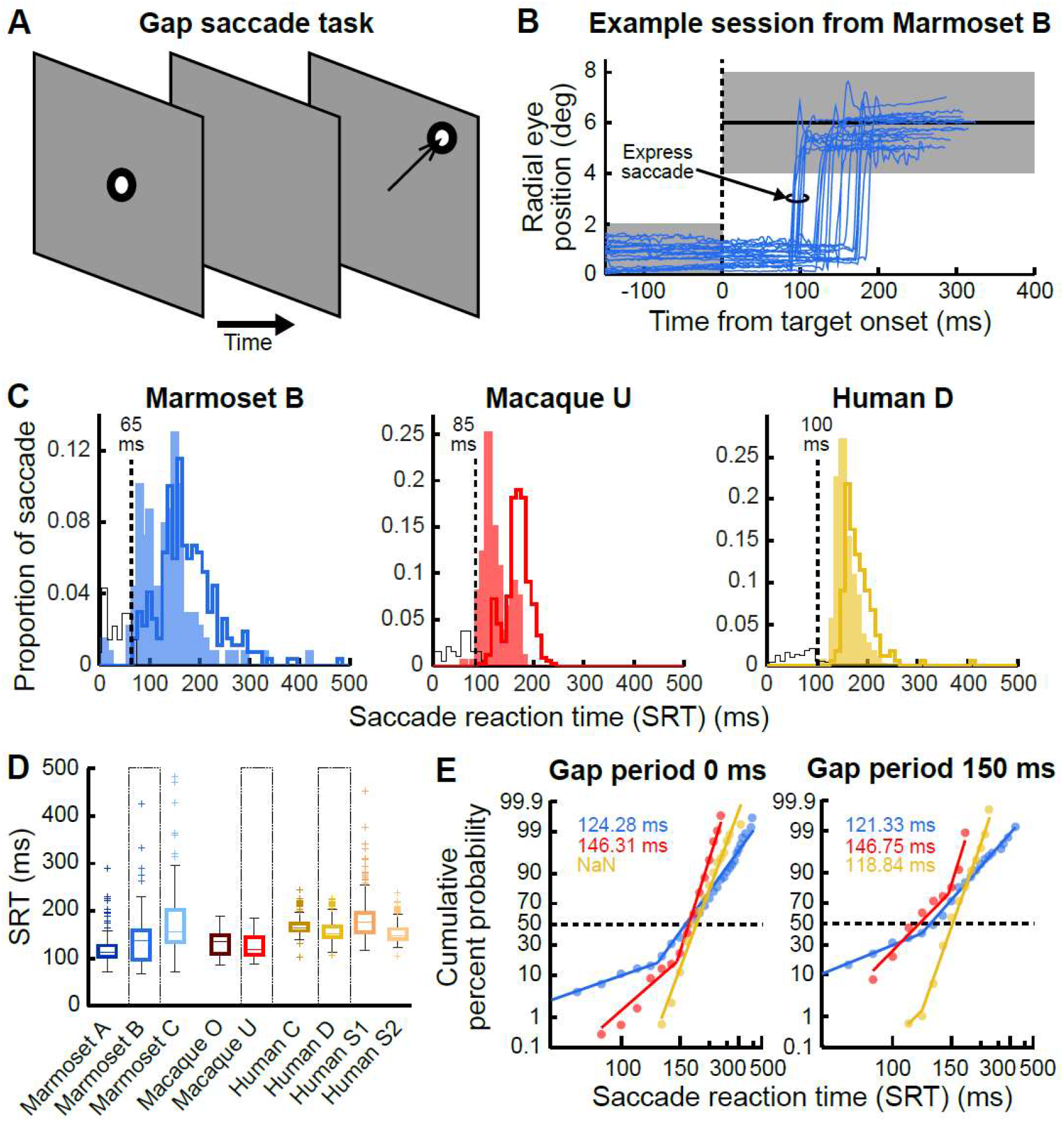
Results of gap saccade task obtained from the three species with 150 ms gap period. **(A)** Diagram of gap saccade task. The convention is the same as Fig. 4A. The subject needed to first fixate at the fixation target for ~200 ms. The fixation spot then disappeared for a certain period of time (gap period), but the subject was required to maintain gaze at the location of the disappeared fixation target during this entire period. The saccade target then appeared and the subject had to saccade to the target to complete one trial. **(B)** Example session from the same subject in Fig. 4B performing gap saccade task with 150 ms gap period. The convention is the same from Fig. 4B. Note that there are short latency saccades made around 100 ms after target onset in this example session which was not observed in Figure 4B. These saccades are marked as express saccades. **(C)** Example SRT histogram for each species. In order to facilitate comparison, the SRT histogram from step saccade task in Fig. 4C is plotted again in the current figure with the same convention. The color shaded area represents the SRT histogram while each example subjects performed gap saccade task with 150 ms gap period. The black hollow histogram indicates the anticipatory saccades (Materials and Methods) for each species. There were enough anticipatory saccades for Marmoset B to show a clear separation between the two distributions. For Macaque U and Human D, the anticipatory saccades were few, so here we plotted all anticipatory saccades collected from all 5 gap periods. The dashed line and the number indicate the threshold for anticipatory saccades for further exclusion in our analysis. **(D)** Box plot for SRT in all subject. Color indicates each subject. The convention is the same as Fig. 4D. **(E)** Reciprobit plot for step saccade (gap period 0 ms) and gap saccade with 150 ms gap period. Color indicates the same example subjects showed in Fig. 5C. The data for each subject is fitted with two piecewise linear regression to find the hinge point. This hinge point is then used as the border for express saccades in each subject individually (Fig. 6). The fitted hinge points for each example subject are colored and indicated in the insets. Note from Human D, the gap period 0 ms has no hinge point, meaning there were no express saccades detected. Also note that the border for each subject (same color for left and right column) is not much different for either Marmoset B or Macaque U.

#### Step and gap saccade task, separating express saccades

The next issue was to define the SRT border between what constitutes an express saccade and what constitutes a regular saccade for each species. To do so, we first plotted the SRTs of all response saccades on a reciprobit plot, following “Carpenter and Williams (1995)”. We then applied two piece-wise linear regression to find the hinge point on the reciprobit plot (Fig. 5E). Because the hinge points for each gap period of a given subject were similar, we took the average from all the gap periods we tested and set the SRT border for each subject. We also calculated the percentage of express saccades versus all saccades using this border (Fig. 6B).

**Figure 6.**
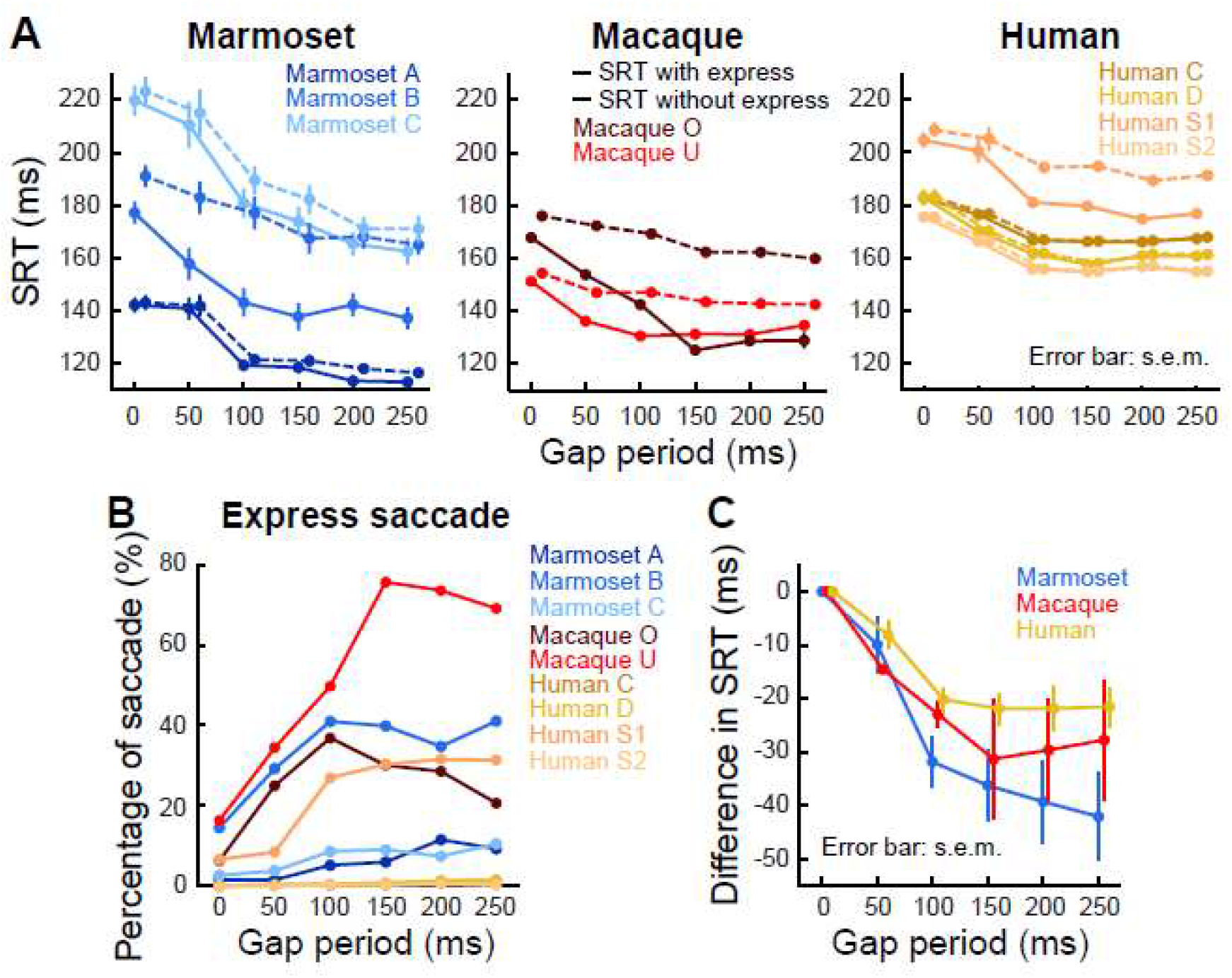
Summary of SRT in gap saccade task. **(A)** Mean SRT from different gap period for all subjects. Each panel plots the results from marmosets, macaques, and humans respectively, from left to right. Colors indicate different subjects. Error bar indicates s.e.m. when visible. **(B)** Percentage of express saccades in each gap period. (**C**) Decrease in mean SRT for each gap period across the three species. The mean SRT of step saccade task (gap period 0 ms in the figure) is used to subtract all the mean SRT in other gap period for each subject. The result is then averaged on a per-species basis. This is to show the change in SRT for each species in gap task. Error bar indicates s.e.m.

#### Statistical analysis

Statistical analysis for all figures is described in each figure legend. Additional statistics performed on the data is reported in the Results or Tables.

### Data and model availability

All data presented in this paper are available upon reasonable request. The source code of the saliency model is published on Github: https://github.com/flyingfalling/salmap_rv. The specific saliency map configuration file used for the analysis is available at: https://github.com/flyingfalling/salmap_rv/models/chenetal2020marmo/params.

## Results

We recorded eye movements while one of the three marmosets, two macaques, or four human subjects watched one of the video clips selected from the same database at original resolution (Fig. 1A). The sampling frequency was 500 Hz for all subjects ensuring the raw eye trace was unbiased among the three species. Using the methods described above, we extracted fixational gaze positions and saccades from the eye movement data. The results are shown for one representative subject from each species in a trial where that subject viewed the same video stimulus (Fig. 1B, C). All three representative subjects from each species share a qualitatively similar distribution of fixations if we simply plot the accumulated fixation time across the whole monitor area, ignoring the time ordering. Furthermore, their saccade trajectories are very similar. Because we did not restrict our subjects’ gaze location, some fixations and saccades fell outside the video, and sometimes even outside the monitor (Fig. 1B, C). However, most fixations fall well within the video size (on average for all subjects all videos separated for each species, marmoset: 92%, macaque: 88%, human: 97% of fixation time fell within the video borders), indicating the video stimuli attracted the subjects’ attention. From the above results, we hypothesized that the three species may share similar bottom-up attention mechanisms which drive their eye movements. We will further elaborate this point using a bottom-up saliency model to predict their gaze location (Fig. 3), which we will present after we characterize the basic saccade kinematics in the next section (Fig. 2).

### Similar saccadic kinematics among the three species in video free-viewing task

After verifying the fixation distribution was similar among marmosets, macaques, and humans, we next compared the basic saccade kinematics of each species. We first looked at the “main sequence”, a stereotypic aspect of saccadic kinematics. When making a saccade, the eye’s peak rotational velocity increases with the saccade amplitude (Bahill et al. 1975). During free-viewing, we observed between 6,000 to 17,000 saccades for each subject of each species, and we individually fit the main sequence for each subject (Table 1). We observed that within each species, subjects’ data ranges were similar (Fig. 2A, 95% of data range overlapped for all subjects in each species). Based on this, for each species we combined all subjects’ saccades into a single set, and calculated main sequence relationship curves on a per-species basis. As shown in Fig. 2A, each species exhibited a stereotypic main sequence. Note that the marmosets were able to make saccades larger than 15 deg, occasionally even making eye movements up to 20 to 30 deg in magnitude while freely viewing the video stimuli. Fig. 2B plots the boundaries of the 95% data range of amplitude against peak velocity for each species. Although the range of saccade amplitude-peak velocity curves overlapped between the three species, the peak velocity tends to be the highest in marmosets, followed by macaques, and finally by humans for a given saccade amplitude.

We next compared inter-saccadic interval for each species. As can be seen in Fig. 2C and Table 2, the median of marmosets was smaller than for macaques and for humans. The inter-saccadic interval for macaques was also shorter than that of humans. From the results of Fig. 2A-C, we also expected the saccade duration for marmosets would be the shortest because the peak velocity was the fastest for a given saccade amplitude. Indeed, marmosets had the shortest duration. We further looked at the saccade amplitude distribution among the three species. We found that human subjects made relatively smaller saccades during the video free-viewing task than the other two species (Fig. 1E, left, Table 2). We reasoned that this could be potentially due to some meaningful features for humans in the videos, e.g. written words which sometimes occur in the video stimuli. Based on this hypothesis, we excluded all saccades from videos that contained structured sentences or phrases from the analysis. Even after excluding videos with sentences or phrases, humans still exhibited a higher proportion of small saccades, visible as a peak on the left side of the graph. (Fig. 1E, right, Table 2). Interestingly, this caused the median of the saccade amplitude significantly decreased for humans (p = 2.86X10^-12^, Wilcoxon rank-sum test). Surprisingly, the saccade amplitude increased for both marmosets (p = 2.05X10^-2^, Wilcoxon rank-sum test) and macaques (p = 8.50X10^-16^, Wilcoxon rank-sum test) in this case. This suggests that small saccades were not just related to reading sentences in the video, but could be derived from the properties of visual search in humans. Our human subjects, for some reason, were more likely to examine local details in the videos than subjects of other species.

**Table 2.** Statistics for saccade kinematics among species. Each row indicating one parameter of the saccade kinematics shown in Fig 2. Each column shows one species comparing to another species. All statistics presented in this table is the p value calculated using Wilcoxon rank-sum test.

|  | Marmoset to<br>Macaque | Macaque to<br>Human | Human to<br>Marmoset |
| --- | --- | --- | --- |
| Inter-saccadic interval | $< 2.23 \times 10^{-308}$ | $3.30 \times 10^{-206}$ | $< 2.23 \times 10^{-308}$ |
| Saccade duration | $< 2.23 \times 10^{-308}$ | $6.88 \times 10^{-34}$ | $< 2.23 \times 10^{-308}$ |
| Saccade amplitude | $< 2.23 \times 10^{-308}$ | $< 2.23 \times 10^{-308}$ | $1.41 \times 10^{-180}$ |
| Saccade amplitude without word | $2.38 \times 10^{-238}$ | $< 2.23 \times 10^{-308}$ | $5.30 \times 10^{-105}$ |

### Similar bottom-up saliency prediction among the three species performing video free-viewing task

To further analyze how the video features attract attention for each species, we implemented the Itti-Koch saliency map model (Itti and Koch 2000). Because the original map was designed for stationary figures and stimuli, we used a modified version which includes motion features (Itti 2005) (Materials and Methods). The saliency map model is a well-established, hypothesis driven, computational model of bottom-up attention which has been proven to have good results in predicting gaze position in both human (Itti 2005) and macaque (White et al. 2017; Yoshida et al. 2012). Examples of saliency map output for one of our video stimuli are shown in Fig. 3A. We quantified the extent to which the saliency map model was able to predict the targets of looking using the area under the receiver operating characteristic (AUROC) for each subject (Fig. 3B-C). Given no assumptions about what a given subject should look at, we were still able to use this bottom-up model to predict the gaze of each subject well above chance (Fig. 3D). Furthermore, the gaze prediction by the model was similar among the three species (AUROC mean per species was 0.60 for marmosets, 0.62 for humans, 0.62 for macaques). Thus, gaze of all species was drawn to bottom-up visual features as predicted by the saliency map model, suggesting that the brains of all species implement a similar baseline bottom-up module that directs attention to visual features that are locally conspicuous.

### Inter-individual difference for visually-guided step saccade task among the three species

To further test if our marmosets could be trained to perform classical saccadic tasks and to compare their performance with macaques and humans, we started with the most basic task: the visually-guided step saccade task (Materials and Methods, Fig. 4A). As shown in Figure 4B, marmosets are capable of being trained to perform this task. During the task, marmosets could accurately fixate and also make single saccades to a peripheral target at 6-degree eccentricity. For comparison, we also had humans and trained macaques perform the task. Fig. 4C-D shows that marmosets tended to have a wider range of saccadic reaction time (SRT) than macaques and humans. To compare the distributions directly between species, we picked one subject from each species with matched SRT medians and looked at their distribution (Fig. 4C, D). The range was the largest for Marmoset B (65-303 ms) after excluding outliers. Macaque U (120-219 ms) and Human D (135-247 ms) were comparable to one another. However, the larger range of SRT was not due to marmosets having slower SRT. The marmosets also had the shortest SRT among the three species (Fig. 4D, Table 3). In other words, both sides of the range were wider. Because we started training all three marmosets on the same day and their training protocol was identical, we expected all marmosets to have mutually comparable SRT distributions. Surprisingly, after six months of training, the median of SRTs from the three marmosets were significantly and noticeably different (Table 3). This result was stable across days. For the data we collected from macaques or even untrained human subjects, the medians, although significant, did not differ at such a magnitude (Table 3).

**Table 3.**
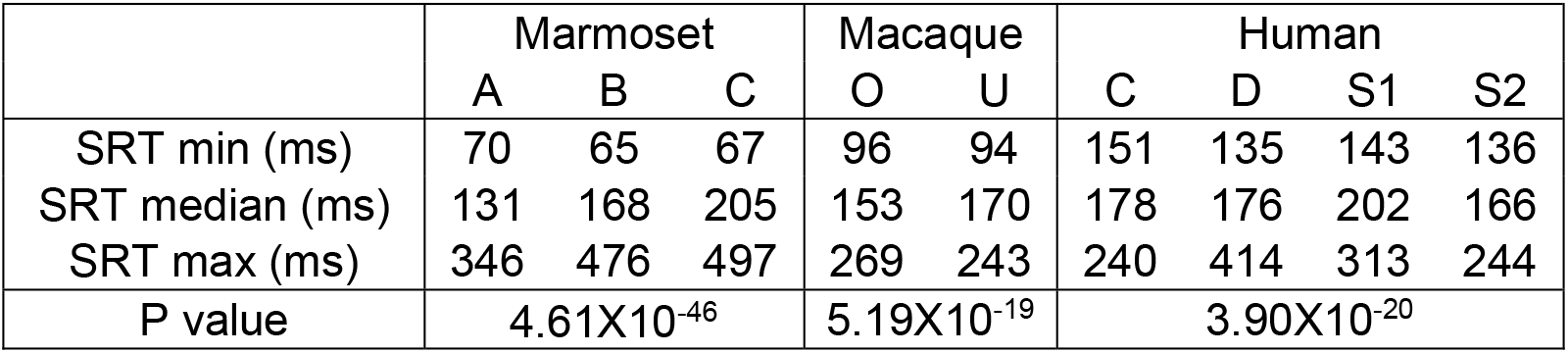
Saccade reaction time (SRT) distribution for step saccade tasks. Each row shows the shortest SRT (SRT min), the median of SRT (SRT median), or the longest SRT (SRT max) for each subject. P value indicates comparing SRT for all subjects of the same species using Kruskal-Wallis test.

|  | Marmoset |  |  | Macaque |  | Human |  |  |  |
| --- | --- | --- | --- | --- | --- | --- | --- | --- | --- |
|  | A | B | C | O | U | C | D | S1 | S2 |
| SRT min (ms) | 70 | 65 | 67 | 96 | 94 | 151 | 135 | 143 | 136 |
| SRT median (ms) | 131 | 168 | 205 | 153 | 170 | 178 | 176 | 202 | 166 |
| SRT max (ms) | 346 | 476 | 497 | 269 | 243 | 240 | 414 | 313 | 244 |
| P value | 4.61X10<sup>-46</sup> |  |  | 5.19X10<sup>-19</sup> |  | 3.90X10<sup>-20</sup> |  |  |  |

### Direction-dependent differences in SRT

We next separated the data by the different target locations and looked at the SRT for each subject (Fig. 4E). This was motivated by previous reports that in macaque (Hafed and Chen 2016; Zhou and King 2002) and human (Abegg et al. 2015), the SRT toward the upper visual field is shorter than that for stimuli in the lower visual field. To verify whether this was the case in our data, we combined the SRT of all target locations in the upper visual field (from positive x axis, 45, 90, 135 deg counter-clockwise) and compared it with the SRT of all visual target locations in the lower visual field (from positive x axis, 225, 270, 315 deg counter-clockwise). For human subjects, three out of four subjects showed significantly shorter SRTs to stimuli in the upper visual field than the lower visual field (Table 4). Also, both macaques showed shorter SRT in the upper visual field. However, in marmosets, only one marmoset (Marmoset A) showed significantly shorter SRT in the upper visual field.

**Table 4.**
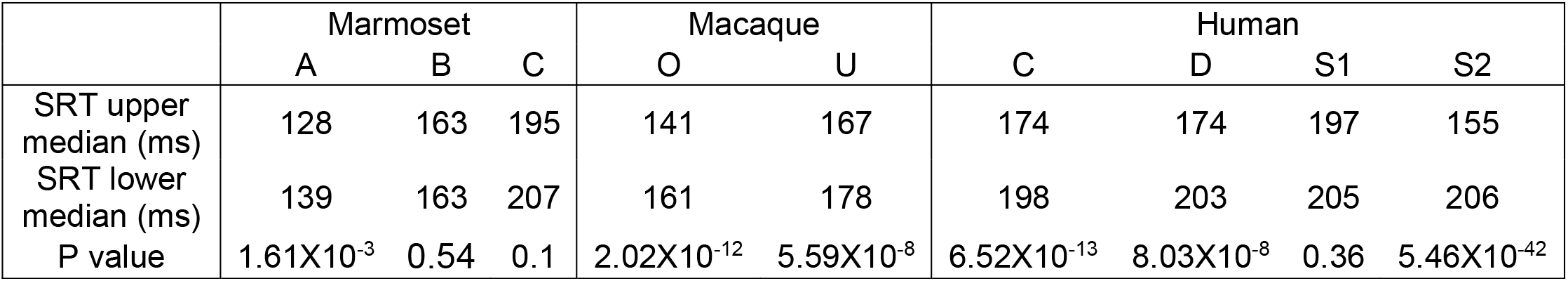
Saccade reaction time (SRT) in upper versus lower visual field for step saccade task. First row shows the SRT median for saccades going to the upper visual field (SRT upper median). Second row shows the SRT median for saccades going to the lower visual field (SRT lower median). P value indicates comparing SRT for upper versus lower visual field for each subject using Wilcoxon rank-sum test.

|  | Marmoset |  |  | Macaque |  | Human |  |  |  |
| --- | --- | --- | --- | --- | --- | --- | --- | --- | --- |
|  | A | B | C | O | U | C | D | S1 | S2 |
| SRT upper median (ms) | 128 | 163 | 195 | 141 | 167 | 174 | 174 | 197 | 155 |
| SRT lower median (ms) | 139 | 163 | 207 | 161 | 178 | 198 | 203 | 205 | 206 |
| P value | $1.61 \times 10^{-3}$ | 0.54 | 0.1 | $2.02 \times 10^{-12}$ | $5.59 \times 10^{-8}$ | $6.52 \times 10^{-13}$ | $8.03 \times 10^{-8}$ | 0.36 | $5.46 \times 10^{-42}$ |

### Decreased SRT and increased express saccade ratio for gap saccade task

After demonstrating that all species can reliably perform basic saccade tasks, our next step was to compare performance in a more sophisticated saccades task: the gap saccade task (Fig. 5A). Fig. 5B-E shows results from trials with a gap period of 150 ms. As demonstrated by data from a single session of marmoset B (Fig. 5B), marmosets are perfectly capable of performing the gap saccade task. There is a clear group of trials in the example session of saccades that have SRTs significantly shorter than the others – these could potentially be express saccades.

To quantify subjects’ performance in the gap saccade task, we overlaid the gap task SRT distributions (Fig. 5C, color shaded area) and the normal step saccade SRT distribution (Fig. 5C, distribution without color shade). The data shown is for the same three subjects used in Fig. 4. It is immediately apparent in the data for marmoset B that there is a set of extremely fast SRTs 0-20 ms after the stimulus onset. These saccades could be anticipatory saccades (Materials and Methods). If they are anticipatory, they are not visually driven and should be excluded (Kalesnykas and Hallett 1987). We separated those trials containing anticipatory saccades and excluded them based on the threshold set for each species (Materials and Methods, Fig. 5C, black outlined distribution for each subject). Under the gap paradigm, the SRT distribution for all subjects shifted to the left, meaning that the SRT was shorter on average for 150 ms gap period than for the step saccade task. Indeed, the median SRT for all subjects is significantly shorter in the gap than in the step saccade task (Table 5).

**Table 5.** Saccade reaction time (SRT) distribution for 150 ms gap period of gap saccade task. Each row shows the shortest SRT (SRT min), the median of SRT (SRT median), or the longest SRT (SRT max) for each subject. P value indicates comparing SRT to step saccade task (Table 3) for each subject using Wilcoxon rank-sum test.

|  | Marmoset |  |  | Macaque |  | Human |  |  |  |
| --- | --- | --- | --- | --- | --- | --- | --- | --- | --- |
|  | A | B | C | O | U | C | D | S1 | S2 |
| SRT min (ms) | 71 | 67 | 71 | 86 | 86 | 102 | 106 | 117 | 105 |
| SRT median (ms) | 113 | 136 | 156 | 135 | 118 | 164 | 153 | 177 | 148 |
| SRT max (ms) | 289 | 424 | 484 | 187 | 183 | 244 | 226 | 453 | 237 |
| P value | 1.13X10<sup>-19</sup> | 1.25X10<sup>-12</sup> | 2.66X10<sup>-12</sup> | 9.29X10<sup>-19</sup> | 1.69X10<sup>-47</sup> | 1.12X10<sup>-38</sup> | 7.01X10<sup>-29</sup> | 2.76X10<sup>-14</sup> | 1.17X10<sup>-38</sup> |

Removal of ostensibly anticipatory saccades yields an SRT distribution that is still clearly bimodal for both marmoset B and macaque U. This indicates that a 150 ms gap period can successfully induce a higher proportion of saccades to be express than a 0 ms gap (i.e. the step saccade task) (Fig. 5C). Using the data from the gap saccade and step saccade task, we delineated the border between what constitutes an express saccade and what constitutes a regular saccade for each subject. To find the border, we calculated the best fit for a two-piece piecewise linear regression on the reciprocal of SRT versus the cumulative probability of that SRT (i.e. a two-piece piecewise reciprobit model). We did this for each subject and for each gap period in our data. The border was then defined as the hinge point between the two pieces of our piecewise regression (Fig. 5E, Materials and Methods). Although the express saccade border varied between subjects, the border did not change much when the gap period was changed (Table 6). We separated SRTs into express saccades and regular saccades using the mean threshold (over all gap periods) from each subject.

**Table 6.**
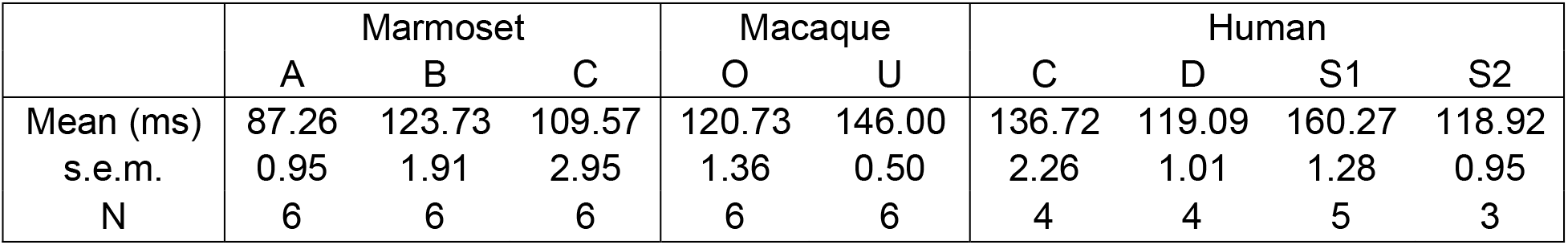
Express saccade boarder from step and gap saccade task. First row shows the mean saccade reaction time (SRT) border calculated with two-piece piecewise linear regression for step saccade task and all gap period gap saccade task and averaged across. Second row shows the standard error of the mean (s.e.m.). Because not all the conditions yielded express saccades, we also indicate with “N” how many of the possible gap periods (0, 50, 100, 150, 200, 250 ms gap period) yielded gap saccades, and thus how many were used to compute the border between express and normal saccades. “N” is used for s.e.m. calculation.

|  | Marmoset |  |  | Macaque |  | Human |  |  |  |
| --- | --- | --- | --- | --- | --- | --- | --- | --- | --- |
|  | A | B | C | O | U | C | D | S1 | S2 |
| Mean (ms) | 87.26 | 123.73 | 109.57 | 120.73 | 146.00 | 136.72 | 119.09 | 160.27 | 118.92 |
| s.e.m. | 0.95 | 1.91 | 2.95 | 1.36 | 0.50 | 2.26 | 1.01 | 1.28 | 0.95 |
| N | 6 | 6 | 6 | 6 | 6 | 4 | 4 | 5 | 3 |

As previously presented, all subjects and all species showed shorter mean SRT under the gap saccade condition than in the step saccade condition (gap period 0 ms) (Fig. 6A, solid line). This is the “gap effect”. This remains true even if we exclude all saccades that we have categorized as express saccades (Fig. 6A, dashed line). We can further also quantify what proportion of a subject’s saccades are express saccades (Fig. 6B). In summary: Marmoset B, Macaque O, Macaque U, and Human S2 have more express saccades than the other subjects. In general, for all subjects, a longer gap period led to a higher proportion of express saccades. Furthermore, most human subjects (Human C, Human D, and Human S2) exhibited almost no saccades that could be classified as express saccades, whereas marmosets and macaques all exhibited express saccades. However, above a gap period of 150 ms, there does not seem to be any higher proportion of express saccades than for shorter gap periods. This means that the shortening in SRT with gap saccade task may have reach a floor effect at gap period 150 ms. This can be clearly visualized by plotting the difference between the mean SRT of each gap period minus the mean SRT for the 0 ms gap (step saccade) condition (Fig. 6C).

## Discussion

We investigated the saccadic behavior of marmosets, macaques, and humans under identical tasks while recording eye movements using similar setups. Under a video free-viewing task, we found that all three species exhibited stereotypic exponential correlation of saccade amplitude to saccade peak velocity, known as the “main sequence.” However, details of their looking behavior, such as the inter-saccadic interval, saccade durations, and saccade amplitudes differed between species. Despite these basic differences, the gaze of all species was drawn to similar locations in the videos at a similar rate. To further support this, we used a model of bottom-up visual attention – the saliency map model – to show that basic visual features such as local differences in color, motion, or luminance draw the attention of all species. After comparing free-viewing behavior, we investigated the similarities between species in convention trained paradigms such as the visually-guided step saccade and gap saccade task. We confirmed that marmosets can be trained on classical saccadic tasks that have been used in macaque and human experiments. We also confirmed that express saccades and the gap effect demonstrated in macaques and humans can also be observed in marmoset data. We conclude that marmoset is a viable alternative animal model for oculomotor and attentional research. The marmoset offers several practical advantages, while not sacrificing the ability to generalize results to human (or macaque, the other traditional model).

Our results on saccade kinematics showed that although all three species exhibited a main sequence in saccades, the actual parameters for both the slope and the exponent were different (Fig. 2A). To the best of our knowledge, this is the first time such a direct comparison among the three species has been reported using such similar experimental conditions. Previous literature has compared marmosets and macaques, and, in contrast to our data, reported the main sequence from both species to be identical (Mitchell et al. 2014). This discrepancy could be due to the previous study having used different eye tracking devices between marmosets and macaques (Chen and Hafed 2013) or due to e.g. different sampling frequency when capturing gaze position (Juhola et al. 1985). To avoid this, we explicitly minimized differences in experimental tasks, data acquisition methods, and data processing between different species. Further support for the veracity of our data lies in its similarity to many previous reports of the main sequence parameter in humans, which has been reported to be 300-400 deg/s for 10 deg saccade (Bahill et al. 1975; Bollen et al. 1993; Port et al. 2016), and previous reports of the peak velocity in macaques, which has been reported to be 500-700 deg/s (Buonocore et al. 2017; Corrigan et al. 2017). These values are repeated by multiple authors, and are comparable with our human and macaque data. Despite the clear difference in parameters between macaque and human that has been reported in previous literature, there has been little discussion of what could be causing this difference, or what its implications are for generalizing results from macaque to human. With data points from a third species, the current study allows us to speculate that primates with smaller body size will have a corresponding increase in saccade peak velocity, given the same saccade amplitude. Several mechanistic explanations could be made for such a trend, such as stronger eye muscles allowing faster contraction after the neural signal arrives or simply smaller eye ball size allowing the muscle to finish contraction faster. Indeed, eye velocity is even faster in smaller animals with smaller eye balls, like mouse (Sakatani and Isa 2007; Wang et al. 2015). Of course, more data points for other primates will be necessary to test this hypothesis.

We used the saliency map model to predict gaze for all three species. In actuality, investigation of the nuances of models that predict eye movements based on visual content is a subject that should deserve a report in its own right. To avoid controversy, we applied a standard baseline model which has been previously used in predicting human gaze (Fig. 3). One justification for this model is that it is inspired by primate early visual cortical processing, which we would argue is largely similar across the three species in this paper (Koch and Ullman 1985).The model has previously been applied to not only human looking behavior, but also macaque (Itti 2005; Itti and Koch 2000; White et al. 2017; Yoshida et al. 2012). More in-depth analysis of why the model predicts looking behavior is necessary – but as a first approximation it indicates that the basic visual information processing mechanisms underlying bottom-up attention in marmoset is similar to macaque and to human. To the best of our knowledge, this is the first time such data is demonstrated among marmoset, macaque, and human. This is also encouraging because this is the first evidence that marmoset can be used not only as a model of oculomotor behavior, but also as a model animal for investigating vision and attention and the neural circuits responsible thereof.

One point of contention that we anticipate is the argument that species of different sizes and shapes and ecological niches will surely have different distributions of neural tunings and sensitivities for spatial frequency, color, and motion in their early visual areas. It could be argued that to predict their behavior using a single model that espouses a single set of tunings is essentially meaningless. In fact, our argument is the opposite – the fact that a single model without specialization is able to predict the gazes of the animals so well is a stronger argument for the similarity of the visual systems of the species. We would also point out that applying a single model to the *same* species may well be meaningless, based on the well-established existence of parallel systems (e.g. magnocellular and parvocellular pathways) within the same species, and even more recent findings that different parts of the visual field may have different tunings within the same individual (Chen et al. 2018)

On that note, we would like to draw attention to a curious aspect observed in the current data. Unlike human and macaque, where saccades to the upper visual field have shorter reaction time than the lower visual field (Abegg et al. 2015; Hafed and Chen 2016; Zhou and King 2002), our marmosets seemed to have less difference in SRT. The origin of this difference could potentially be ecological in nature, relating to constrains in the daily life of each species, which pressured different functional specialization between upper and lower visual fields (Previc 1990). From this point of view, a primate that does not stand upright on the ground for most for their lifespan, but rather (like marmoset) tends to hang upside down in trees, would have no functional advantage of specialization between upper and lower visual fields. The marmoset could be used to test this hypothesis, since even in cavity, the marmoset hang upside down often. Indeed, our data suggested such asymmetry between upper and lower visual field, at least at the level of SRT, was not noticeable (Fig. 4E). We believe this interesting property of marmoset will further raise research on comparing neuronal functions between upper and lower visual fields in macaque and marmoset, and work to reveal the evolutionary changes across different primates.

It has been shown that in both human and macaque, saccade reaction times are shortened under gap paradigms (Saslow 1967). This has been termed the gap effect, and can even induce ultra-fast, minimum latency saccades called express saccades (Fischer and Boch 1983; Fischer and Ramsperger 1984). It has since been shown that the occurrence of express saccades is dependent on experimental conditions such as the length of gap period, training, and target predictability (Carpenter 2001; Paré and Munoz 1996). In the current study, we demonstrated conclusively that marmosets not only exhibit a gap effect (Johnston et al. 2018) but with proper training, can also be induced to make express saccades, just like humans and macaques (Fig. 5-6) One possible confound to this conclusion is that we did not randomize the initial fixation period, and thus made the task simpler for the marmosets. The marmosets were able to time the delay between the upcoming gap period and saccade target. We also observed that the SRT of marmoset express saccades were shorter than other species, but this is unsurprising since marmoset reaction times are shorter even under the basic visually-guided step saccade task (Fig. 4). Several studies indicated the generation of express saccades could be involved in “disinhibition of fixation” and preparation of the upcoming saccade through several potential cortical or subcortical brain areas (Aizawa et al. 1999; Chen et al. 2013; Dias and Bruce 1994; Everling and Munoz 2000; Isa and Hall 2009; Paré and Munoz 1996; Schiller et al. 2008; Schiller and Tehovnik 2005). However, how these brain areas interact with one another has not been directly demonstrated. One potential approach to understand such complex neural interaction would be to use large scale, even brain-wide neural recording with high time and spatial resolution (Isa 2017; Tia et al. 2017; Umeda et al. 2019). The high resolution necessitated by the short time gaps from one saccade to the next. Such high-accuracy measurement is difficult in macaque because many critical target brain areas lie in the deep sulci in macaque. Targeting one area with high precision is already difficult if it is in a sulcus, to say nothing of simultaneous recording from multiple areas. Marmoset, however, has a lissencephalic brain (smooth) brain, and so is not limited by these technical difficulties. This, combined with other advantages such as its small body size, moves us to contend that the marmoset as a model animal will soon help us advance our understanding of the mechanism behind express saccades.

Although all individual marmosets were trained in comparable amount of time, their results in classical saccadic tasks differed significantly, indicating differences in personality or ability. For example, in the visually-guided step saccade task, the median of SRT for marmosets had the largest difference, even larger than the largest difference if we allow comparisons between macaques and humans (Fig. 4D). We do not know the reason for such large inter-individual variance, which we could not correct even with extensive training. However, it is important to note that even with such a large difference between individuals, all conclusions related to the basic saccade kinematics (Fig. 2), saliency prediction (Fig. 3), upper and lower saccade latency (Fig. 4), gap effect, and also express saccade (Fig. 5-6) hold true. Our results are highly consistent and similar to results observed in macaques and humans. In addition to reporting and directly comparing the oculomotor characteristics of marmoset with macaque and human, this study also provides copious evidence that the marmoset is a suitable animal model for research in oculomotor and visual attention control.

## Acknowledgements

We thank Yusuke Yamamoto and Norihiro Takakuwa for assisting us gathering data from macaques. Chih-Yang Chen was a JSPS International Research Fellow. The current study was funded by the Brains and Minds Project No. 18dm0207020h0005 to T.I., Project No. 19dm0207093h0001 to H.O., Project No. JP19dm0207069 to M.Y. and K.M. from AMED, Japan, and by Grant-in-Aid for Scientific Research (#20K06920) to KM from JSPS, Japan.

## References

Abegg M, Pianezzi D, Barton JJS. A vertical asymmetry in saccades. J Eye Mov Res 8: 1–10, 2015.

Aizawa H, Kobayashi Y, Yamamoto M, Isa T. Injection of nicotine into the superior colliculus facilitates occurrence of express saccades in monkeys. J Neurophysiol 82: 1642–1646, 1999.

Bahill AT, Clark MR, Stark L. The main sequence, a tool for studying human eye movements. Math Biosci 24: 191–204, 1975.

Belmonte JCI, Callaway EM, Caddick SJ, Churchland P, Feng G, Homanics GE, Lee K-F, Leopold DA, Miller CT, Mitchell JF, Mitalipov S, Moutri AR, Movshon JA, Okano H, Reynolds JH, Ringach DL, Sejnowski TJ, Silva AC, Strick PL, Wu J, Zhang F. Brains, Genes, and Primates. Neuron 86: 617–631, 2015.

Bollen E, Bax J, van Dijk JG, Koning M, Bos JE, Kramer CGS, van der Velde EA. Variability of the main sequence. Invest Ophthalmol Vis Sci 34: 3700–4, 1993.

Borji A, Tavakoli HR, Sihite DN, Itti L. Analysis of Scores, Datasets, and Models in Visual Saliency Prediction. In: 2013 IEEE International Conference on Computer Vision. IEEE, p. 921–928.

Buonocore A, Chen C-Y, Tian X, Idrees S, Münch TA, Hafed ZM. Alteration of the microsaccadic velocity-amplitude main sequence relationship after visual transients: implications for models of saccade control. J Neurophysiol 117: 1894–1910, 2017.

Carpenter RHS. Express saccades: is bimodality a result of the order of stimulus presentation? Vision Res 41: 1145–1151, 2001.

Carpenter RHS, Williams MLL. Neural computation of log likelihood in control of saccadic eye movements. Nature 377: 59–62, 1995.

Chen C-Y, Hafed ZM. Postmicrosaccadic enhancement of slow eye movements. J Neurosci 33: 5375–5386, 2013.

Chen C-Y, Sonnenberg L, Weller S, Witschel T, Hafed ZM. Spatial frequency sensitivity in macaque midbrain. Nat Commun 9: 2852, 2018.

Chen M, Liu Y, Wei L, Zhang M. Parietal Cortical Neuronal Activity Is Selective for Express Saccades. J Neurosci 33: 814–823, 2013.

Corrigan BW, Gulli RA, Doucet G, Martinez-Trujillo JC. Characterizing eye movement behaviors and kinematics of non-human primates during virtual navigation tasks. J Vis 17: 1–22, 2017.

Dias EC, Bruce CJ. Physiological correlate of fixation disengagement in the primate’s frontal eye field. J Neurophysiol 72: 2532–2537, 1994.

Everling S, Munoz DP. Neuronal correlates for preparatory set associated with pro-saccades and anti-saccades in the primate frontal eye field. J Neurosci 20: 387–400, 2000.

Felleman DJ, Van Essen DC. Distributed Hierarchical Processing in the Primate Cerebral Cortex. Cereb Cortex 1: 1–47, 1991.

Fischer B, Boch R. Saccadic eye movements after extremely short reaction times in the monkey. Brain Res 260: 21–26, 1983.

Fischer B, Ramsperger E. Human express saccades: extremely short reaction times of goal directed eye movements. Exp Brain Res 57: 191–195, 1984.

Fuchs AF. Saccadic and smooth pursuit eye movements in the monkey. J Physiol 191: 609–631, 1967.

Hafed ZM, Chen C-Y. Sharper, Stronger, Faster Upper Visual Field Representation in Primate Superior Colliculus. Curr Biol 26: 1647–1658, 2016.

Hafed ZM, Krauzlis RJ. Microsaccadic Suppression of Visual Bursts in the Primate Superior Colliculus. J Neurosci 30: 9542–9547, 2010.

Homman-Ludiye J, Bourne JA. The marmoset: An emerging model to unravel the evolution and development of the primate neocortex. Dev Neurobiol 77: 263–272, 2017.

Hubel DH, Wiesel TN. Anatomical Demonstration of Columns in the Monkey Striate Cortex. Nature 221: 747–750, 1969.

Isa T. Intrinsic processing in the mammalian superior colliculus. Curr Opin Neurobiol 12: 668–677, 2002.

Isa T. Using the common marmoset for neurophysiological studies of neocortical functions. J Physiol 595: 7013–7013, 2017.

Isa T, Hall WC. Exploring the superior colliculus in vitro. J Neurophysiol 102: 2581–2593, 2009.

Itti L. Quantifying the contribution of low-level saliency to human eye movements in dynamic scenes. Vis cogn 12: 1093–1123, 2005.

Itti L, Carmi R. Eye-tracking data from human volunteers watching complex video stimuli. CRCNS.org 2009.

Itti L, Koch C. A saliency-based search mechanism for overt and covert shifts of visual attention. Vision Res 40: 1489–1506, 2000.

Johnston KD, Barker K, Schaeffer L, Schaeffer D, Everling S. Methods for chair restraint and training of the common marmoset on oculomotor tasks. J Neurophysiol 119: 1636–1646, 2018.

Juhola M, Jäntti V, Pyykkö I. Effect of sampling frequencies on computation of the maximum velocity of saccadic eye movements. Biol Cybern 53: 67–72, 1985.

Kalesnykas RP, Hallett PE. The differentiation of visually guided and anticipatory saccades in gap and overlap paradigms. Exp Brain Res 68: 115–121, 1987.

Knöll J, Pillow JW, Huk AC. Lawful tracking of visual motion in humans, macaques, and marmosets in a naturalistic, continuous, and untrained behavioral context. Proc Natl Acad Sci U S A 115: E10486–E10494, 2018.

Koch C, Ullman S. Shifts in selective visual attention: towards the underlying neural circuitry. Hum Neurobiol 4: 219–27, 1985.

Krauzlis RJ. The control of voluntary eye movements: New perspectives. Neuroscientist 11: 124–137, 2005.

Kümmerer M, Wallis TSA, Bethge M. Saliency Benchmarking Made Easy: Separating Models, Maps and Metrics. In: Computer Vision – ECCV 2018. Lecture Notes in Computer Science. Springer International Publishing, p. 798–814.

Markov NT, Vezoli J, Chameau P, Falchier A, Quilodran R, Huissoud C, Lamy C, Misery P, Giroud P, Ullman S, Barone P, Dehay C, Knoblauch K, Kennedy H. Anatomy of hierarchy: Feedforward and feedback pathways in macaque visual cortex. J Comp Neurol 522: 225–259, 2014.

May PJ. The mammalian superior colliculus: laminar structure and connections. Prog Brain Res 151: 321–78, 2006.

Medendorp WP, Buchholz VN, Van Der Werf J, Leoné FTM. Parietofrontal circuits in goal-oriented behaviour. Eur J Neurosci 33: 2017–2027, 2011.

Mitchell JF, Leopold DA. The marmoset monkey as a model for visual neuroscience. Neurosci Res 93: 20–46, 2015.

Mitchell JF, Reynolds JH, Miller CT. Active Vision in Marmosets: A Model System for Visual Neuroscience. J Neurosci 34: 1183–1194, 2014.

Moschovakis AK, Scudder CA, Highstein SM. The microscopic anatomy and physiology of the mammalian saccadic system. Prog Neurobiol 50: 133–254, 1996.

Mustari MJ. Nonhuman primate studies to advance vision science and prevent blindness. ILAR J 58: 216–225, 2017.

Paré M, Munoz DP. Saccadic reaction time in the monkey: advanced preparation of oculomotor programs is primarily responsible for express saccade occurrence. J Neurophysiol 76: 3666–3681, 1996.

Park JE, Zhang XF, Choi S-H, Okahara J, Sasaki E, Silva AC. Generation of transgenic marmosets expressing genetically encoded calcium indicators. Sci Rep 6: 34931, 2016.

Port NL, Trimberger J, Hitzeman S, Redick B, Beckerman S. Micro and regular saccades across the lifespan during a visual search of “Where’s Waldo” puzzles. Vision Res 118: 144–157, 2016.

Previc FH. Functional specialization in the lower and upper visual fields in humans: Its ecological origins and neurophysiological implications. Behav Brain Sci 13: 519–542, 1990.

Sakatani T, Isa T. Quantitative analysis of spontaneous saccade-like rapid eye movements in C57BL/6 mice. Neurosci Res 58: 324–331, 2007.

Sasaki E, Suemizu H, Shimada A, Hanazawa K, Oiwa R, Kamioka M, Tomioka I, Sotomaru Y, Hirakawa R, Eto T, Shiozawa S, Maeda T, Ito M, Ito R, Kito C, Yagihashi C, Kawai K, Miyoshi H, Tanioka Y, Tamaoki N, Habu S, Okano H, Nomura T. Generation of transgenic non-human primates with germline transmission. Nature 459: 523–527, 2009.

Saslow MG. Latency for Saccadic Eye Movement. J Opt Soc Am 57: 1030, 1967.

Schall JD. Visuomotor Functions in the Frontal Lobe. Annu Rev Vis Sci 1: 469–498, 2015.

Schall JD, Purcell BA, Heitz RP, Logan GD, Palmeri TJ. Neural mechanisms of saccade target selection: Gated accumulator model of the visual-motor cascade. Eur J Neurosci 33: 1991–2002, 2011.

Schiller PH, Kendal GL, Slocum WM, Tehovnik EJ. Conditions that alter saccadic eye movement latencies and affect target choice to visual stimuli and to electrical stimulation of area V1 in the monkey. Vis Neurosci 25: 661–673, 2008.

Schiller PH, Tehovnik EJ. Neural mechanisms underlying target selection with saccadic eye movements. Prog Brain Res 149: 157–171, 2005.

Shinoda Y, Takahashi M, Sugiuchi Y. Brainstem neural circuits for fixation and generation of saccadic eye movements. Prog Brain Res 249: 95–104, 2019.

Solomon SG, Rosa MGP. A simpler primate brain: The visual system of the marmoset monkey. Front Neural Circuits 8: 1–24, 2014.

Tatler BW, Baddeley RJ, Gilchrist ID. Visual correlates of fixation selection: effects of scale and time. Vision Res 45: 643–659, 2005.

Tia B, Takemi M, Kosugi A, Castagnola E, Ansaldo A, Nakamura T, Ricci D, Ushiba J, Fadiga L, Iriki A. Cortical control of object-specific grasp relies on adjustments of both activity and effective connectivity: a common marmoset study. J Physiol 595: 7203–7221, 2017.

Tomioka I, Ishibashi H, Minakawa EN, Motohashi HH, Takayama O, Saito Y, Popiel HA, Puentes S, Owari K, Nakatani T, Nogami N, Yamamoto K, Noguchi S, Yonekawa T, Tanaka Y, Fujita N, Suzuki H, Kikuchi H, Aizawa S, Nagano S, Yamada D, Nishino I, Ichinohe N, Wada K, Kohsaka S, Nagai Y, Seki K. Transgenic Monkey Model of the Polyglutamine Diseases Recapitulating Progressive Neurological Symptoms. eNeuro 4: ENEURO.0250-16.2017, 2017a.

Tomioka I, Nogami N, Nakatani T, Owari K, Fujita N, Motohashi H, Takayama O, Takae K, Nagai Y, Seki K. Generation of transgenic marmosets using a tetracyclin-inducible transgene expression system as a neurodegenerative disease model. Biol Reprod 97: 772–780, 2017b.

Umeda T, Koizumi M, Katakai Y, Saito R, Seki K. Decoding of muscle activity from the sensorimotor cortex in freely behaving monkeys. Neuroimage 197: 512–526, 2019.

Vanni S, Hokkanen H, Werner F, Angelucci A. Anatomy and Physiology of Macaque Visual Cortical Areas V1, V2, and V5/MT: Bases for Biologically Realistic Models. Cereb Cortex 1–35, 2020.

Veale R, Hafed ZM, Yoshida M. How is visual salience computed in the brain? Insights from behaviour, neurobiology and modelling. Philos Trans R Soc Lond B Biol Sci 372: 20160113, 2017.

Walther D, Koch C. Modeling attention to salient proto-objects. Neural Networks 19: 1395–1407, 2006.

Wang L, Liu M, Segraves MA, Cang J. Visual Experience Is Required for the Development of Eye Movement Maps in the Mouse Superior Colliculus. J Neurosci 35: 12281–12286, 2015.

White BJ, Berg DJ, Kan JY, Marino RA, Itti L, Munoz DP. Superior colliculus neurons encode a visual saliency map during free viewing of natural dynamic video. Nat Commun 8: 14263, 2017.

Yarbus AL. Eye Movements and Vision. Plenum Press (New York), 1967.

Yoshida M, Itti L, Berg DJ, Ikeda T, Kato R, Takaura K, White BJ, Munoz DP, Isa T. Residual Attention Guidance in Blindsight Monkeys Watching Complex Natural Scenes. Curr Biol 22: 1429–1434, 2012.

Zhang L, Tong MH, Marks TK, Shan H, Cottrell GW. SUN: A Bayesian framework for saliency using natural statistics. J Vis 8: 32, 2008.

Zhou W, King W. Attentional sensitivity and asymmetries of vertical saccade generation in monkey. Vision Res 42: 771–779, 2002.

